# High temperature limits on developmental canalization in the ascidian *Ciona intestinalis*

**DOI:** 10.1101/488668

**Authors:** Steven Q. Irvine, Katherine B. McNulty, Evelyn M. Siler, Rose E. Jacobson

## Abstract

The normal embryogenesis of marine animals is typically confined to a species-specific range of temperatures. Within that temperature range development results in a consistent, or canalized, phenotype, whereas above and below the range abnormal phenotypes are produced. This study reveals an abrupt high temperature limit, occurring over a 1-2°C range, for normal embryonic development in *C. intestinalis*. Above that threshold morphological abnormalities in the notochord and other organs are observed, beginning with cleavage and gastrula stages, and becoming more pronounced as embryogenesis proceeds. However, even in highly morphologically abnormal temperature disrupted (TD) embryos, cell type specification, including muscle, endoderm, notochord, and sensory pigment cells is accomplished. An explanation for this finding is that in *C. intestinalis* cell type specification occurs relatively early in embryogenesis, due to cleavage stage segregation of maternal cytoplasmic determinants and short-range cell interactions, which are largely intact in TD embryos. On the other hand, morphogenesis of the notochord and other structures is dependent on precise cell movement and shape changes after the gastrula stage, which appear to be disrupted above the high temperature threshold. These findings have implications for the relationship between ecology and reproduction in *C. intestinalis*. More broadly they point to mechanisms behind canalization in animals, such as ascidians, characterized by early, largely autonomous, cell type specification.

## Introduction

What is the weak link in embryogenesis that limits developmental success at temperature maxima? This question is central to understanding the limits of an organism’s ecological niche, for which temperature is a major determinant. This study seeks to establish which aspects of embryogenesis are the most susceptible to high temperature in the model marine invertebrate *Ciona*.

*C. intestinalis*, like many if not all animals, has a “normal” embryonic phenotype that is produced over a range of environmental conditions. The ability for a developmental program to produce a stereotyped outcome in spite of environmental variation has been termed “canalization” (Siegal and Bergman, 2002; Waddington, 1942).

While there has been much work done on the effects of water temperature on the overall life history of marine invertebrates, a literature search reveals little work examining in detail water temperature effects on embryogenesis itself. However, it is likely that the ability to develop to a functional larval stage is an imporftant factor limiting the ranges of many marine invertebrates (Byrne et al., 2009). It is also likely that embryologists historically have been more interested in how normal development operates than in how it goes awry due to environmental conditions (Gilbert, 2011). On the other hand, perturbation of normal processes, experimentally or by mutation, are standard experimental approaches to understanding normal developmental processes (Gilbert, 2006). The study of perturbation of normal development by excessive temperature could make use of findings from experimental embryology as clues to the critical links in maintenance of the phenotype.

Temperature is a major determinant of marine invertebrate ranges (e.g. (Bhaud et al., 1995; Pörtner and Gutt, 2016), life history distribution (Marshall et al., 2012), community ecology (Dijkstra et al., 2011), and developmental mode (Olive, 1995; Pappalardo and Fernandez, 2014). Notably, changes in range have been shown to track changes in water temperature (Poloczanska et al., 2013). There are various aspects of life history that could be affected by temperature. One major focus of ecological research has been on the effects of temperature on physiological performance in adult animals (e.g. Bolton et al., 2013; Evans and Hofmann, 2012; 2013; Somero, 2011). However development, before the larval stage, is sensitive to environmental temperature, which may be limiting in certain cases (Byrne, 2011). This would be true especially in animals that have a long breeding period and that may see a wide range of temperatures, such as *C. intestinalis* in southern New England waters. Given these factors an open area of research would be the proximate physiological and embryological mechanisms that are potentially disrupted by temperature anomalies. If these mechanisms can be identified the data would point to aspects of the phenotype subject to selection under new temperature regimes.

The genus *Ciona*, including the morphologically nearly indistinguishable *C. intestinalis* (formerly *C. intestinalis*, sp. B), and *C. robusta* (*C. intestinalis*, formerly sp. A), has abundant scientific resources in developmental genetics and genomics that could be brought to bear once the temperature-related critical points in development are identified (Lemaire, 2011; Satoh, 1994; Satoh and Levine, 2005; Stolfi and Christiaen, 2012). These resources will provide background to suggest approaches for identifying the molecular developmental mechanisms that are disrupted by temperatures outside the “normal” range.

This paper describes experiments to determine the temperature maxima for canalized normal development in *C. intestinalis*. Development to the larval stage was examined to identify when and how embryogenesis is disrupted at the temperature maximum. It was found that different cell types, such as notochord and pigmented ocellus are present in embryos deformed due to high temperature. However, those cells fail to find their proper positions and morphology during embryogenesis. These findings suggest that processes leading to the specification of cell type are robust to high temperature in *C. intestinalis*, but that mechanisms involving cell behaviors, such as movement and differential adhesion, are perturbed by temperatures above a critical range.

## Materials and Methods

### Animal collection

Adult *C. intestinalis* (formerly *C. intestinalis*, sp. B) were obtained from floating docks in the Point Judith Marina in Point Judith Pond in southern Rhode Island, USA (41.387°N, 71.517°W). Animals were maintained in closed aquaria at 15-18°C under constant light until use. Field sea surface temperatures at the collection site were recorded using HOBO UA-002-08 data loggers (Onset Computer Corp.).

### Temperature experiments

Gametes were obtained by dissection of two different adults for each brood. Eggs were maintained at 18°C on a chilled platform during washing with 0.4 μm filtered seawater (FSW) and subsequent steps. Dechorionation was done chemically, using 0.4 mg/ml Pronase E (Sigma P5147) in 1% sodium thioglycolate (w/v) in FSW, at 18°C and placed at the respective incubation temperatures in environmental chambers to equilibrate for 1-4 hours (see text). Eggs were then fertilized with mixed sperm from two adults, resulting in mixed broods of sibling and half-sibling embryos. Embryos were fixed at various time points using either 4% formaldehyde in PTw (1x PBS; 0.1% Tween-20), or 2% paraformaldehyde in Ca/Mg-free seawater (CMFSW: NaCl 463 mM; KCl 11 mM; Na_2_SO_4_ 25.5 mM; NaHCO_3_ 2.1 mM; HEPES, pH 8.0 10 mM) for 30 minutes. After fixation embryos were washed 4 times with PTw.

Some experiments were done with undechorionated embryos as noted. These were scored after hatching. They were fertilized as for the dechorionated embryos after the eggs equilibrated to the incubation temperatures. All percentages are the means of the mean score for individual broods per temperature and time point. Refer to Fig. 1 legend for statistics and information on numbers of broods and embryos scored. Developmental stage numbers, where noted, refer to the scheme in Hotta et al. (2007).

**Figure 1.**
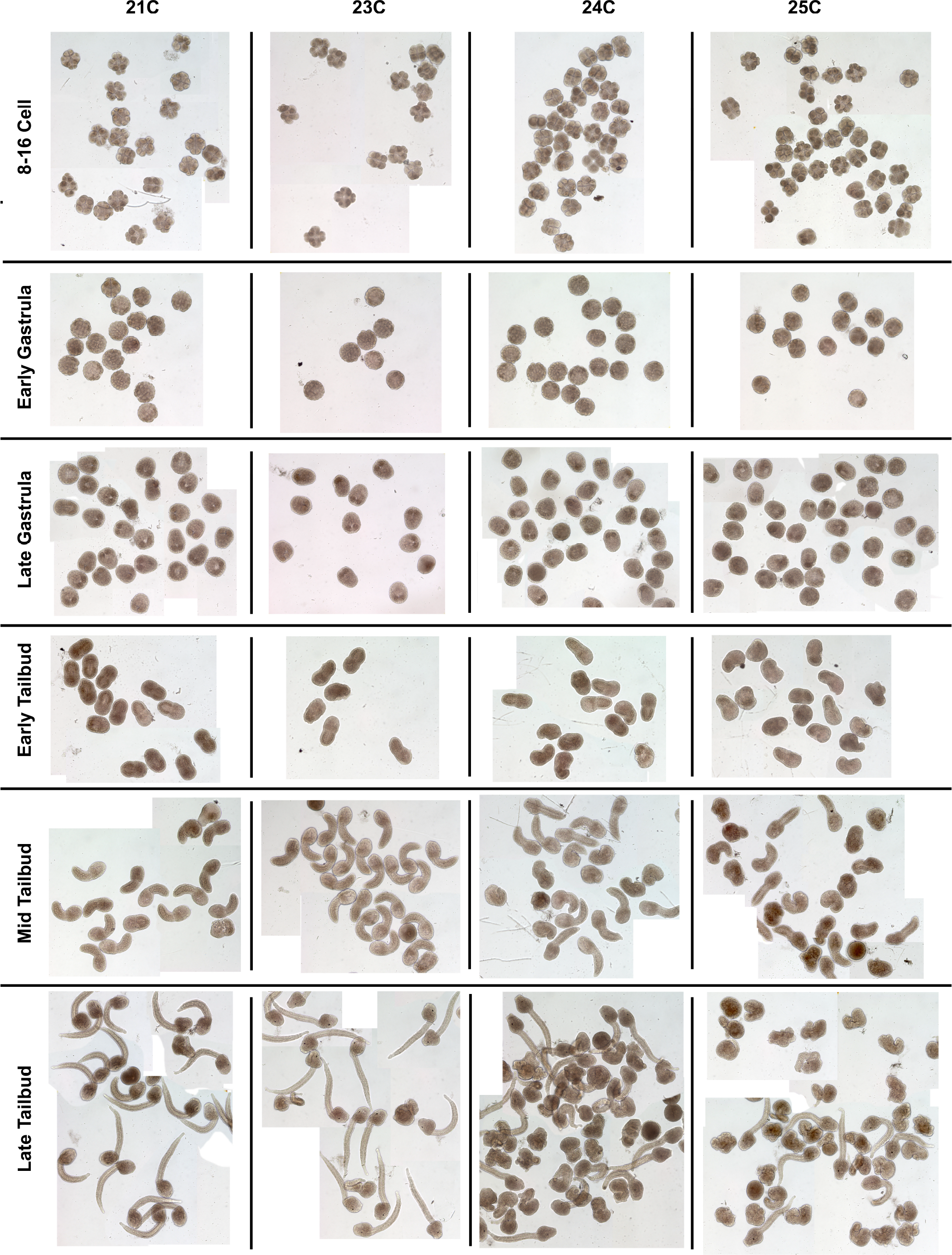
A) Sea surface temperatures at the collection site measured by electronic data loggers. Red arrows indicate collection dates for animals used as parents for embryos that provided data included in this study. B) Mean percentages of normal larvae from half sibling broods produced at high temperatures during June - July 2018. 3-5 independent broods scored at each temperature with an average of 134 embryos scored per brood per temperature (min. 48, max. 310). Error bars are +/− one standard deviation of the means. ** 23° and 24°C means are significantly different at p<0.01 by t-test. C) Percentages of various larval phenotypes (ref. Table 1) from the same half sibling broods as in 1B. Error bars and numbers of experiments and larvae scored as in 1B. D) Mean percentages of normal development for early stage embryos (up to 6 hpf) from independent dechorionated half sibling broods. Parent animals collected during the same time period as for B and C above. The apparent increase in normal development at 25°C from 4-5 h. is likely a scoring artifact. 3-5 broods scored at each temperature and time point with a median of 58 total embryos scored per temperature/time (min. 48, max. 101). Error bars are +/− one standard deviation of the means. *** p<0.001 for difference between the means by 2-tailed t-test. Other comparisons were not significantly different.

### Histochemistry and phalloidin staining

Alkaline phosphatase (AP) and acetylcholinesterase staining was done as described in Swalla (2004). F-actin and nuclei were visualized similar to Christiaen et al. (2005): embryos were fixed in 2% paraformaldehyde in CMFSW for 30 min.; permeabilized with PBST2 (1x PBS; 0.2% Triton X-100) plus 50 mM ammonium chloride for 30 minutes at room temperature; stained with 0.4 U Alexa Fluor 546-phalloidin (Molecular Probes cat. no. A22283) and 5μg/ml DAPI (4′,6-diamidino-2-phenylindole) in PBST2 for 2 hours at room temperature with gentle agitation in the dark; then washed once with PBST1 (1x PBS; 0.01% Triton X-100) for 5 minutes and twice with PBS for 10 minutes. Specimens were mounted in glycerol: PBS 1:1 w/v with 1% DABCO for confocal imaging.

## Results

### Rearing temperatures and developmental success

To observe morphological effects of high seawater temperatures on development, one sibling/half-sibling cohort of dechorionated embryos were reared at 4 temperatures, 21-25°C, and samples photographed at various stages (Fig. 1; chorions were removed because morphology is not clearly seen in embryos inside their chorions.). The proportion of cleavage stage anomalies can be seen to increase at the higher temperatures. Likewise, later stage disruptions also increase, with an abrupt drop in the proportion of normal embryos in the 1°C difference between 23°C and 24°C (Fig. 2B-D).

**Figure 2.**
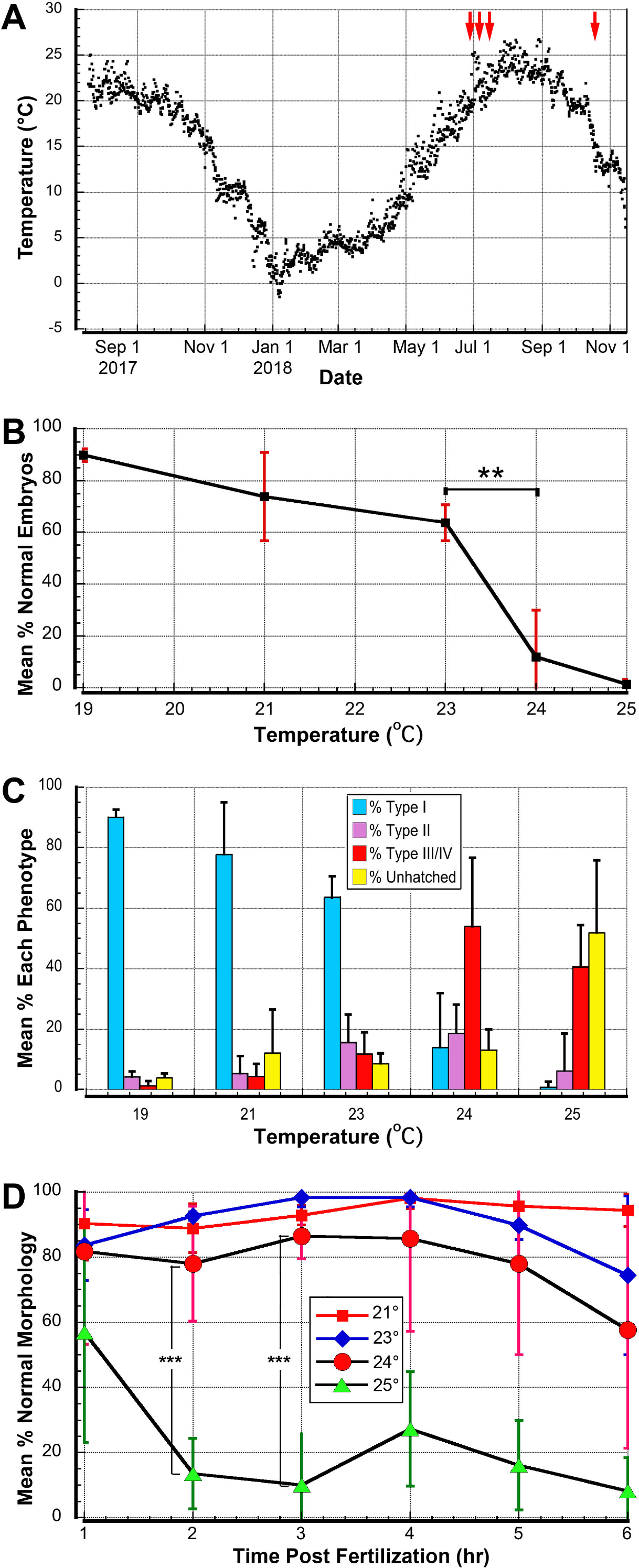
A single half sibling brood reared at different temperatures in June 2018 and photographed at similar stages. Nearly all larvae appear normal through 23°C but the percentage of normal larvae falls of steeply at 24°C, and nearly all larvae are abnormal at 25°C. Interestingly, the early stages through gastrula appear nearly normal at all temperatures (although ref. Fig. 5 for details of abnormalities at early stages).

At our estuarine collection site in Rhode Island, USA, water temperature varied in 2017-2018 from a mid-summer high of 26.7°C to a mid-winter low of −1.4°C (Fig. 2A). The diurnal temperature variation is typically around 2°C in summer. Parent animals used in this study were collected during two time periods: June 24-July 10, 2018, when the water temperature varied between 18.0 and 25.3°C, and October 21, 2018, when the water temperature was 14.1-15.0°C (red arrows in Fig. 2A). During August and September, 2018 we failed to obtain high percentages of normal embryos at any tested incubation temperature, even 18°C, after dechorionation. The chemical dechorionation process is likely quite stressful, but high percentages of dechorionated eggs from animals collected at lower field water temperatures develop normally.

To quantify the effects of water temperatures at the high end of the local range, we scored the developmental success of embryos from parents collected June-July 2018, incubated at 19-25°C. For this set of experiments, the unfertilized eggs were equilibrated at the incubation temperature for one hour before fertilization. These were scored for normal morphology at the larval stage after hatching. We found that the percentage of normally developed larvae fell gradually up to 23°C, then dropped significantly between 23°C and 24°C (Fig. 2B, p<0.01, 2-tailed t-test).

The embryonic phenotypes resulting in the high temperature rearing experiments were scored as 4 types (Table 1): Type 1, “control”, typical of embryos reared at permissive temperatures; Type II, “kinked tail”; Type III, “deformed tail”, and Type IV, “globular”. Fig. 2C shows the distribution of larval phenotypes from the same experiments as Fig. 2B, including those embryos that fail to hatch. At 25°C many embryos failed hatching, possibly due to a mechanical inability to break open the chorion, or an arrest of development prior to hatching stage. Again, note a large increase in abnormal temperature disruption (TD) phenotypes above 23°C.

**Table 1.** Phenotype definitions

| Phenotype | Nickname | Specific late stage characteristics |
| --- | --- | --- |
| Type I | normal | Typical wild type late tailbud: tail - straight notochord, 20 notochord cells, straight lateral muscle bands, dorsal nerve cord, ventral endodermal strand; trunk - 3 palps, lateral atrial rudiments, dorsal sensory vesicle with 2 pigment cells, etc. |
| Type II | kinked tail | Trunk similar to Type I; tail with 20 notochord cells but shorter due to partial failure of extension, and with abnormal kink(s) |
| Type III | deformed tail | Failure to progress to normal late stage trunk morphology; notochord cells fail to fully extend, resulting in much shorter curled tail |
| Type IV | globular | Highly anomalous trunk morphology, including abnormal position of pigment cell(s), lack of clear sensory vesicle, abnormal overall shape; complete failure of extension of notochord cells result in disorganized tail anatomy |

**Table 2.**
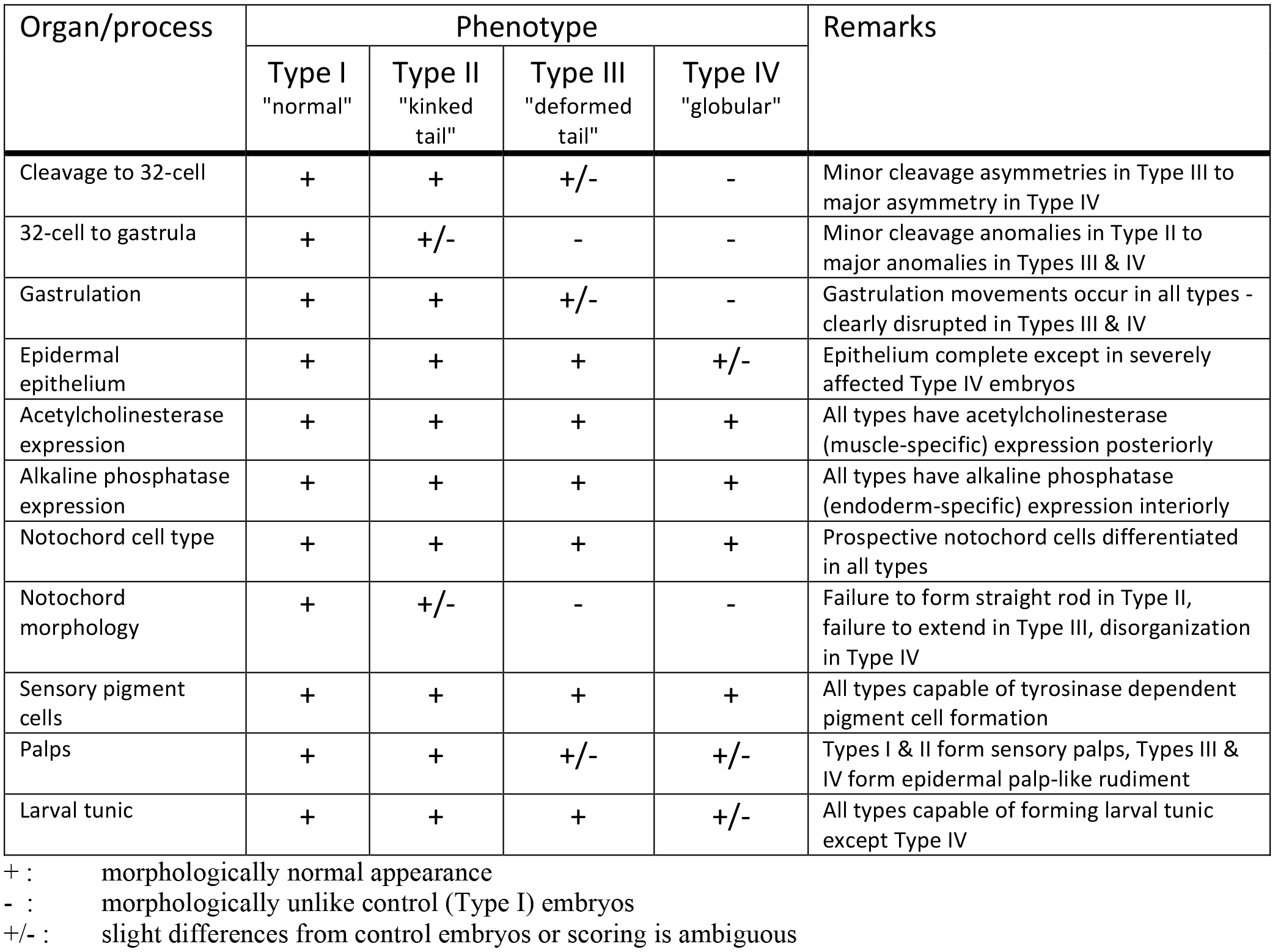
Phenotype scoring of TD embryos

To examine whether early stages are susceptible to TD we fixed subsets of dechorionated broods at 1 hr. intervals and scored them for normal cleavage stage morphology (Fig. 2D). Through 6 hrs. of development minor deviations from normal morphology were observed in broods at 21 - 24°C. At 25°C nearly all embryos from 2 hrs. on exhibited highly abnormal cleavage. Developmental rate is positively related to temperature, so at 25°C the embryos reach the 32-cell - 64-cell stages by 3 hrs., as compared with 16-cells at 18°C. Mean percentages of normal morphology were significantly different (2-tailed t-test) between 24 and 25°C at the 2 and 3 hr. time points. These data indicate that, at least up to 3 hours, development is resistant to TD relative to later stages, since embryos reared at 24°C, which will result in a high percentage of abnormal larvae, have largely morphologically normal early cleavage patterns. Because of high variance within the high temperature broods at other time points the differences between temperature conditions are not statistically significant.

### Effects on cell type specification

It was observed that even in batches of high temperature embryos in which a very high proportion of extremely deformed larvae are produced, most of them express pigment spots in the two melanocytes located in the sensory vesicle (Figs. 1 & 3). The melanin pigment in these cells does not appear until late in embryogenesis (Stages 23-25; Hotta, 2007) suggesting that the melanocyte cell type specification pathway, is not disrupted throughout development. To test whether other cell type specification pathways were intact or disrupted in TD embryos, histochemical assays for other cell type-specific markers were performed.

**Figure 3.**
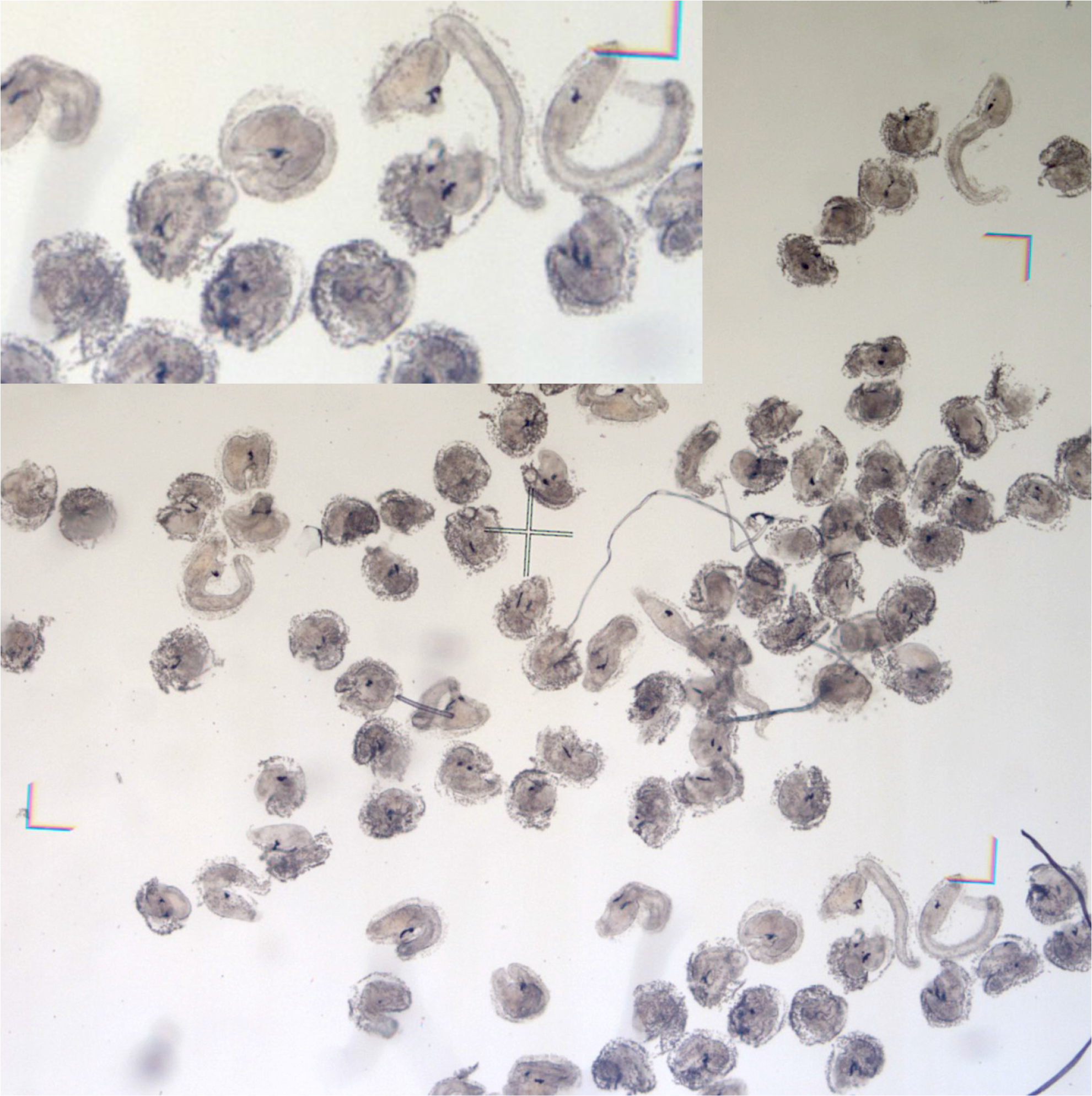
A group of TD half sibling larvae. Note that even severely disrupted larvae have pigment cells dependent on late stage expression of melanin.

In control early and late tailbud embryos alkaline phosphatase (AP) histochemistry resulted in endodermal staining in the trunk, as expected (Fig 4A,B). Similarly, AP expression is apparent in the trunks of TD embryos (Fig. 4C-E). In testing for muscle cell specification, acetylcholinesterase (AChE) histochemistry was positive in tail muscle of controls (Fig. 4F,G). Staining in TD embryos at the same stages corresponds with that of the controls (Fig. 4H-J). These assays indicate that basic endoderm and muscle cell type specification is accomplished, even in Type IV TD embryos.

**Figure 4.**
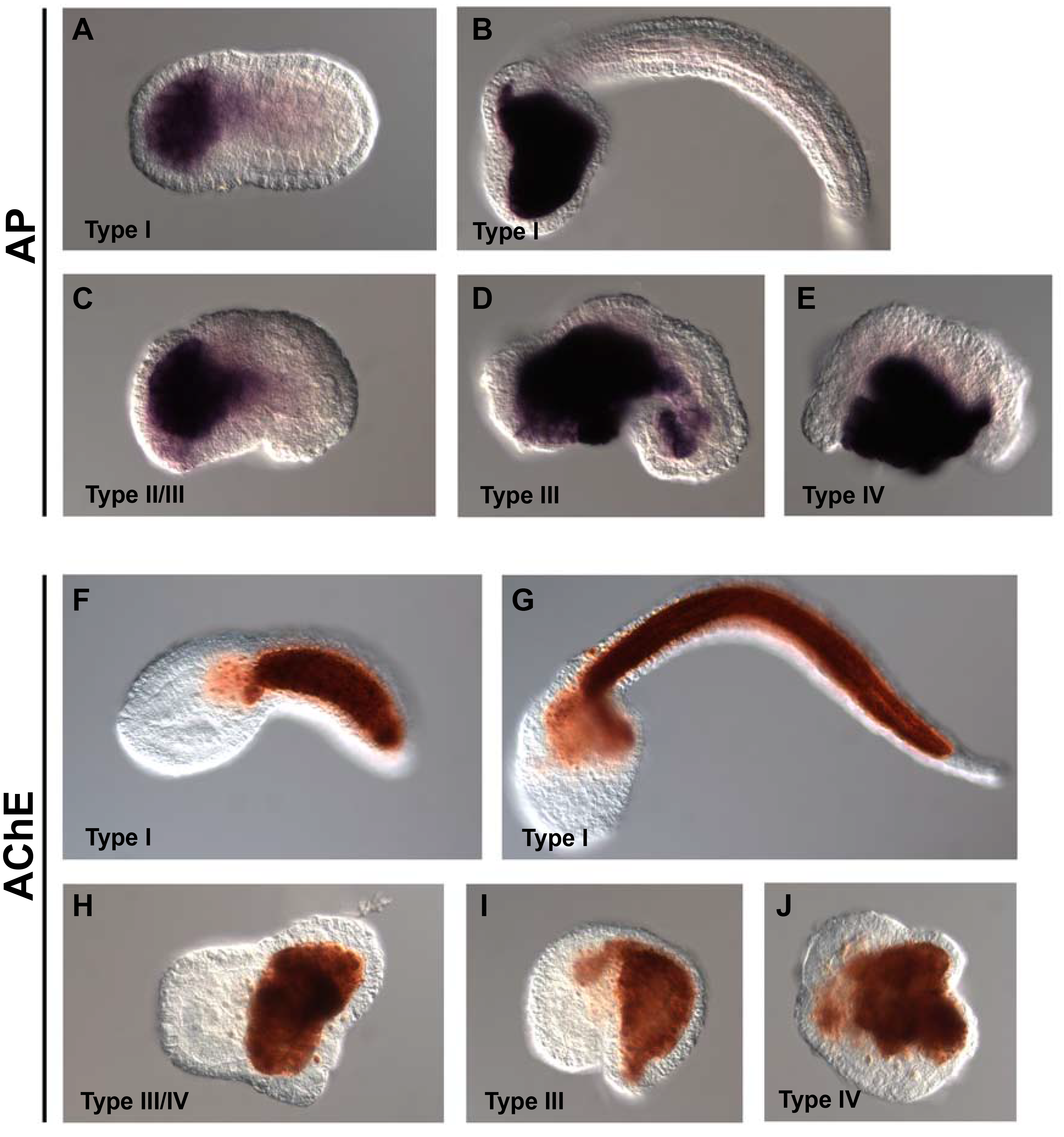
Histochemical assays for alkaline phosphatase (A-E) and acetylcholinesterase (F-J) protein expression, indicative of endoderm and muscle respectively. A) Initial tailbud and B) late tailbud Type I (normal) embryos. C) Initial tailbud TD embryo. D, E) late tailbud TD embryos. F) mid tailbud and G) late tailbud Type I embryos. H) mid tailbud, and I, J) late tailbud TD embryos.

### Effects on embryonic morphology

In order to determine at which embryonic stages effects of high temperature appear, individual embryos incubated at those temperatures were followed through embryogenesis and photographed at roughly corresponding stages (Fig. 5). As suggested by Fig. 1, in Type II TD embryos (Fig. 5B-C) slight asymmetries and other anomalies appear by the gastrula stage which become more pronounced in Type III and Type IV embryos (Fig. 5D-G). These patterns contrast with the clearly symmetrical and stereotypical morphology of a Type I embryo (Fig. 5A). It is important to note that these embryos were reared from parents collected in late October, when seawater temperatures were lower than for the experiments described above. Similar levels of TD were observed at 22°C in these broods as were only obtained at 24°C for the summer experiments.

**Figure 5.**
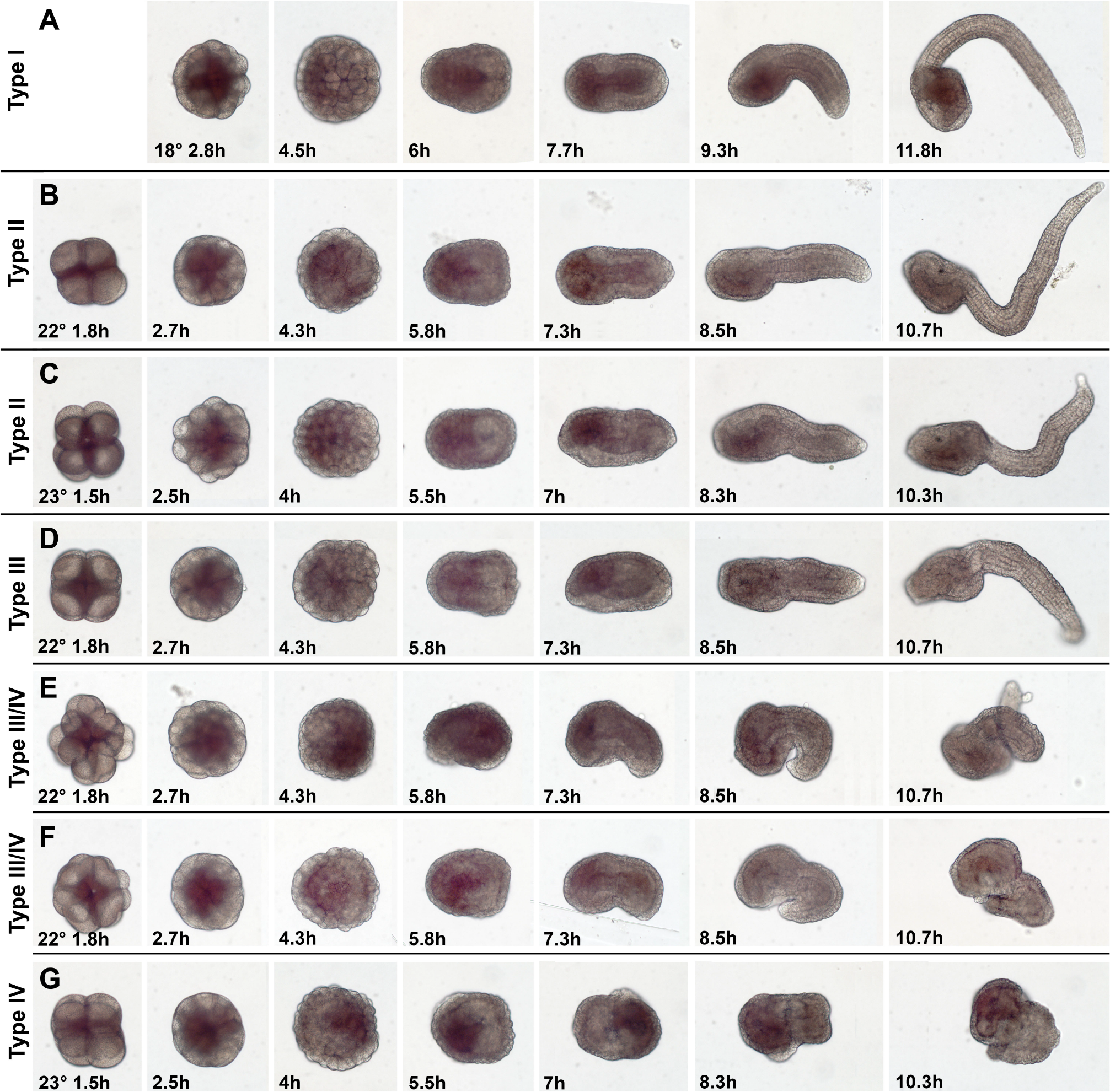
Individual embryos photographed through embryogenesis. All embryos from the same half sibling brood pre-incubated for 3 hours at the rearing temperature before fertilization. The parents for this brood were collected October 21, 2018 when field seawater temperatures were lower than for the experiments conducted in June - July (Fig. 1A). Note that morphological abnormalities are observed at lower temperatures in this brood than in those of parents collected in the warmer summer water (cf. Fig. 2). Rearing temperatures and time post fertilization as listed. More severely affected (Type III - IV) embryos exhibit more apparent abnormalities in cleavage and gastrulation than Type II embryos, indicating that these early anomalies contribute to the severity of the temperature disruption. All embryos, however, differentiate pigment cells at the time equivalent of the late tailbud stage.

Temperatures above threshold can produce highly abnormal cleavage patterns (cf. Fig. 6A & B). More severely affected embryos, however, may not persist to gastrula stage. Fig. 6D & E show TD gastrulae which exhibit anomalous cell shapes as compared with a control gastrula (cf. Fig. 6C’-E’, arrowed and double arrowhead regions). TD embryos at all tested temperatures invariably still make an epidermal single layered epithelium (Fig. 6G/G’ & I/I’, open arrowheads). All embryos examined also make notochord cells, as judged by their large cell size and interior position. However, control embryos (Fig. F/F’ & H/H’), have an intercalating column of cells with a smooth curved overall shape. TD embryos, on the other hand, have notochords that range from a kinked column (Fig. 6G/G’, solid arrowhead) to disorganized columns (Fig. 6I/I’) to masses of large cells that fail to assemble in a column at all (Fig. 7D’, arrow).

**Figure 6.**
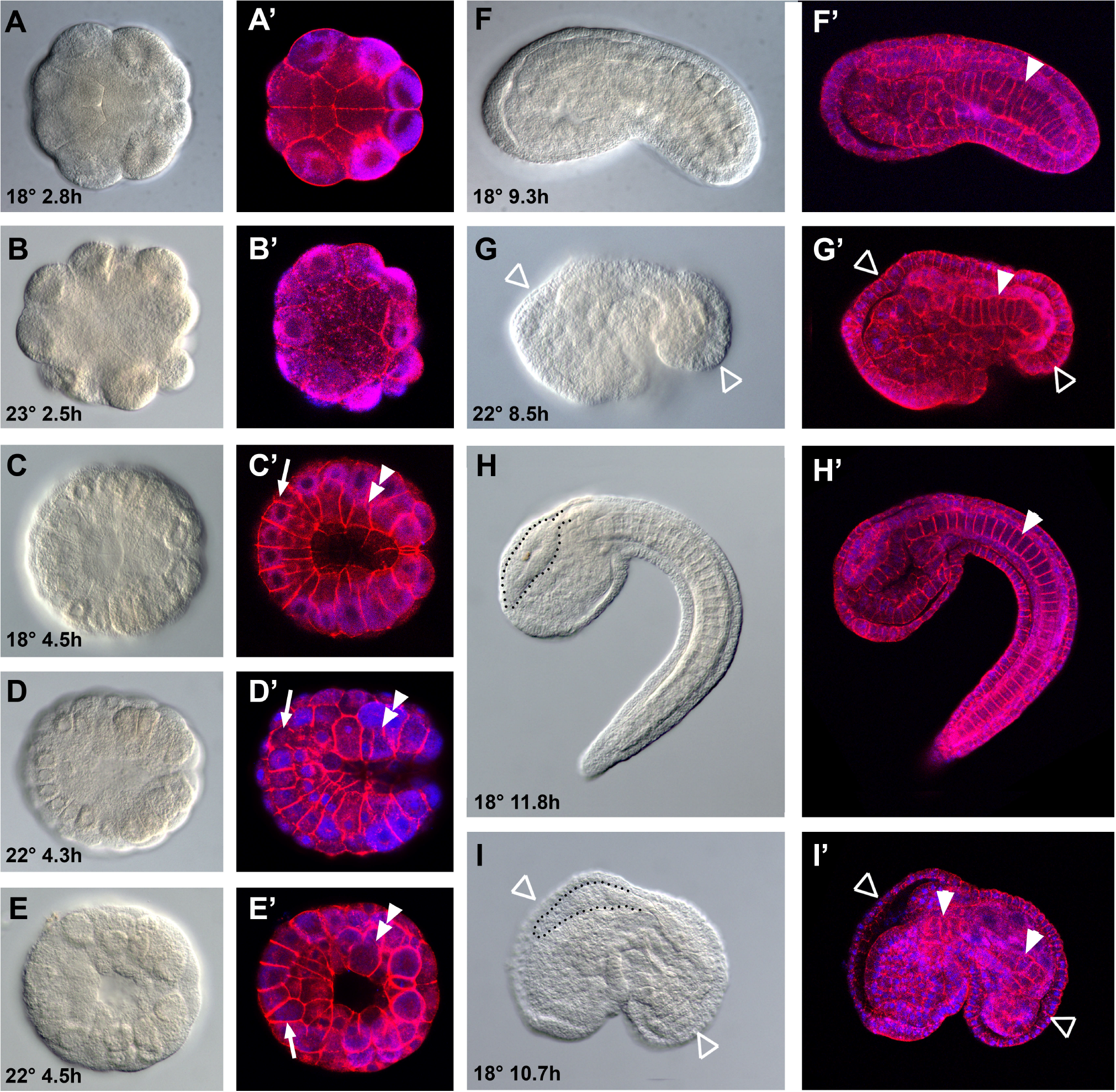
Representative normal (Type I; A, C, F, H) and TD (Type III/IV; B, D, E, G, I) embryos at cleavage to mid gastrula stages, from same brood as in Fig. 5. Each embryo is depicted in DIC (left) and phalloidin/DAPI (right) pairs, with rearing temperature and TPF as listed.

**Figure 7.**
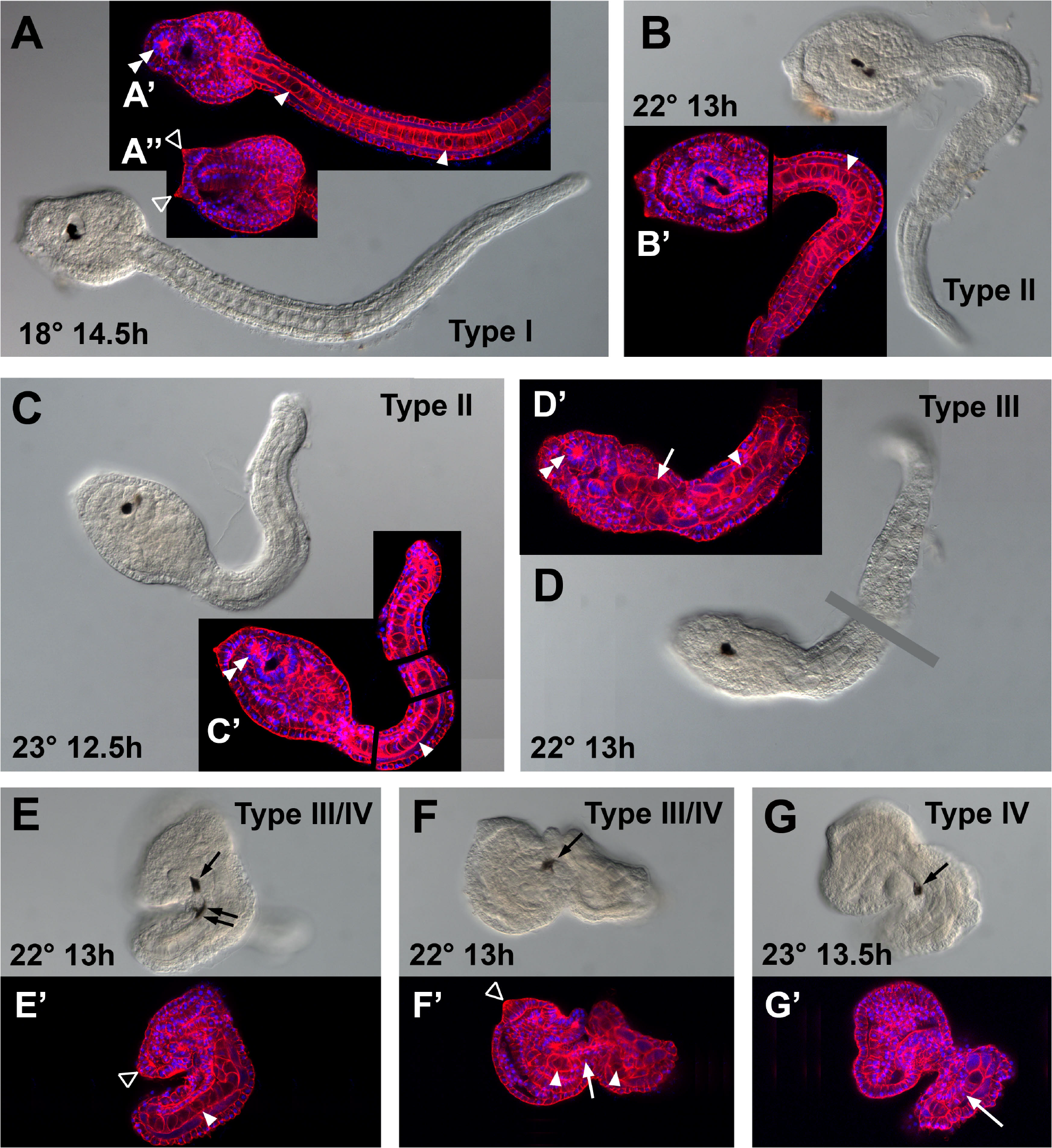
Higher magnification views of larvae of individuals from Fig. 5. DIC and phalloidin/DAPI images are paired, with rearing temperature and TPF as listed. Letters denoting individual animals correspond to those in Fig. 5 (e.g., A, A’, A”, photographs are of the same animal denoted A in Fig. 5). Arrowheads, vacuolated notochord cells; double arrowheads, stomodeum; open arrowheads, sensory palps; white arrows, abnormal notochord morphology; black arrow, sensory pigment cell; double black arrow, putative differentiated ocellus pigment cell.

The same individual embryos depicted in Fig. 5 were examined in more detail as late tailbuds (Stage 24-25) using DIC microscopy and phalloidin/DAPI staining (Fig. 7). In Type II embryos (Fig. 7B/B’, C/C’) the trunk appears morphologically normal, including the formation of a sensory vesicle and stomodeum (double arrowheads in Fig. 7A’, C’). The notochord cells form a single file and become vacuolated (Fig. 7B’, C’; single arrowheads), as in control embryos (Fig. 7A’). However, the tails fail to extend properly and do not assume the smooth curve seen in controls. In Type III TD embryos (Fig. 7D/D’, E/E’) the notochord cells sometimes fail to intercalate properly (Fig. 7D’, arrow), and in other cases intercalate to form a single file but are severely kinked (Fig. 7E/E’). In Type IV embryos the notochord cells may form a single file (Fig. 7F’, G’, arrows) and the cells may become vacuolated (Fig. 7E’,F’, arrowheads) but the notochord extension fails, resulting in lack of tail outgrowth.

As noted previously, melanized pigment cells form even in Type IV embryos (Fig. 7F, G, arrows). However, in the disorganized trunks of Type III and IV embryos the cells are often found in anomalous positions, such as the posterior locations in Fig. 7E and 7G]. Many TD embryos form two sensory cells, and in some cases it is apparent that differentiation between the two cell types, otolith and ocellus has occurred (e.g. Fig. 7E, single vs. double arrows).

Pyramidal anterior epidermal thickenings are present in some severely abnormal Type III and IV embryos (Fig. 7E’ & F’, open arrowheads) suggesting that they are able to form at least rudimentary sensory palps. These structures are characterized by columnar cells in the anterior epidermis which form at Stage 24 (Fig. 7A”, open arrowheads). The presence of the epidermal thickening indicates that regionalization of the epidermis to locate the palp is accomplished in spite of TD. More detailed work would be required to determine if these palp-like structures are fully developed and functional palps.

## Discussion

### Developmental canalization

The finding that normal embryogenesis is maintained over a wide temperature range (i.e. is canalized), but falls off sharply near the maximum local temperature suggests that this canalization is an adaptation to the local temperature regime (cf. Figs. 1 & 2). Anecdotally, *C. intestinalis* from more northern locations have a lower temperature maximum for normal development (T. Meedel, personal communication) indicating that different populations are locally adapted with respect to temperature. *Ciona* sp. has also been reported from tropical locations (Dybern, 1965), with water temperatures higher than the maximum found in our study. It may be that the tropical populations breed in the winter, and not in summer, as is the case for *C. robusta* in the Mediterranean (Caputi et al., 2015). If that is the case, and given that both *C. robusta* and *C. intestinalis* are generally described as temperate or cold water species, there may be some intrinsic constraint to adaptation for development at temperatures higher than about 25°C.

It is also likely that within a population the temperature maximum is plastic depending on the temperature profile experienced by the adults during gametogenesis. In fact, we found that embryos derived from parents collected in late October 2018 had an approximately 2°C lower high temperature threshold than those from parents collected in June-July 2018, although precise quantification of this seasonal plasticity was beyond the scope of this study. A similar seasonal difference in the optimal range for embryogenesis was observed for *C. savignyi* in Japanese waters near Tokyo (Nomaguchi et al., 1997).

In terms of cellular stress and temperature maxima, Sato et al. (2015) have shown that locally sympatric *C. intestinalis* and *C. robusta* have different, genetically programmed, high temperature thresholds for normal development. That study focused on cellular temperature stress, showing that endoplasmic reticulum related chaperones have different expression profiles in the different species. We did not assay for cell stress in this study, but our previous work showed significant upregulation of stress-related proteins, such as Hsc71 and glutathione peroxidase, in adult ovaries of animals from the same population acclimated to 22°C *vs.* those at 18°C (Lopez et al., 2017). Based on the above, it can be assumed that embryos are experiencing a cellular stress response at temperatures approaching the developmental maximum.

One aspect of the experimental design that should be pointed out, is that for these experiments gametogenesis occurred in adults acclimated to temperatures at or below the control temperature of 18°C, while the eggs used were equilibrated to the experimental temperatures for a relatively short time of 1-3 hours before fertilization. It is likely that these eggs mount a heat shock response due to the rapid temperature change, which could affect embryogenesis. On the other hand, in our experience once temperatures in the field reach the upper range tested here, embryos from gametes of parents acclimated at field temperatures exhibit poor development, even when reared at control temperatures. Therefore, it is likely that the temperature disruption mechanisms behind the defects seen in the laboratory temperature conditions, are similar to those operating in nature.

### Temperature disruption and cell type specification

Histochemical assays show that expression of enzymes characteristic of muscle and endoderm is present in TD embryos (Fig. 4). These cell types are well known to be autonomously specified by early segregation of maternal cytoplasmic determinants (Satoh, 1994; Whittaker, 1973; Whittaker et al., 1977; Whittaker and Meedel, 1989). Since cell-cell interactions are not required for the basic specification of these cell types, even in the case of abnormal cleavage patterns one would expect that some proportion of cells would express these markers. In all cases the cells expressing either acetylcholinesterase (muscle) or alkaline phosphatase (endoderm) form contiguous masses, indicating that the cell lineages leading to the segregation of cytoplasmic constituents did not break up into different regions of the embryo.

One of the first observations in our experiments was that melanization associated with the ocellus and otolith, which occurs long after gastrulation, appears even in severely deformed TD embryos (Fig. 3). These differentiation events occur very late in development, with melanization of the second of the two pigment cells appearing only just before the larval stage. Much is known about the cell type specification pathways for these cells (Esposito et al., 2015). They are derived from the bilateral a8.25 cells and require induction from vegetal blastomeres at the 32-cell stage for later pigment expression (Nishida, 1987; Nishida and Satoh, 1989; Oonuma et al., 2016). It has been shown that BMP/Chordin signaling is required in the neural plate for pigment cell formation (Darras and Nishida, 2001a; Miya et al., 1997). In addition, both FGF and Wnt signaling occurring at the tailbud stage is involved in differentiation of the pigment cells (Racioppi et al., 2014; Squarzoni et al., 2011). Hence, at least to the tailbud (Stage 18) cell-cell relationships are intact enough to allow BMP, FGF, and Wnt signaling events to occur. However, it is not known whether the cells expressing pigment always follow the exact lineage of those in control embryos, since multiple cells are competent to become pigment cells (Nishida and Satoh, 1989).

Another aspect of sensory cell development is the differentiation of the otolith vs. ocellus pigment cells. The bilateral a8.25 precursor cells are an equivalence group and apparently become differentiated according to their anterior-posterior position by Wnt signaling (Abitua et al., 2012; Nishida and Satoh, 1989). This differentiation is present at least in some TD cases, such as the embryo in Fig. 7E, which has a more heavily pigmented anterior cell, indicative of the otolith (single arrow), and crescent shaped posterior cell similar to a normal ocellus (double arrow).

Notochord cells are produced in all TD embryos at the temperatures we tested, as judged by their characteristic morphology and position (Figs. 6 & 7). Differentiation of notochord from other mesodermal derivatives requires induction from other vegetal blastomeres prior to gastrulation by FGF and BMP signals (Darras and Nishida, 2001b; Nakatani and Nishida, 1994). While not all vegetal blastomeres are capable of inducing notochord, the induction is not exclusive to the normal inducers, indicating that even if cell-cell relationships are not completely normal, the inductive signal could still be present (Nakatani and Nishida, 1994). At least Type II and III TD embryos have notochord cells capable of becoming vacuolated, which occurs in the late tailbud (Stage 28) in normal Type I embryos. Once specified, isolated notochord cells can go on to vacuolate, so proper cell-cell interactions are not required for that aspect of notochord cell development (Munro and Odell, 2002).

Epidermis is derived from animal hemisphere blastomeres, and all TD embryos in our tests produced a single layered epithelium morphologically similar to that of the epidermis of controls (cf. Fig. 7A/A’ and 7G/G’). Reverberi (1971) showed that isolated a4.2 cell quarter embryos make an epidermal epithelium, so this aspect of epidermal specification is already established early in cleavage (Satoh, 1994). Therefore, the formation of a complete epidermal epithelium may be an intrinsic property of prospective epidermis, remaining operative in abnormal TD embryos.

The sensory palps are derived from ectodermal lineages and are visible as groups of columnar cells in the epithelium of late tailbud embryos (Fig. 7A”, open arrowhead). Type II and III TD embryos are capable of forming palp-like structures in what appears to be the anterior trunk epidermis (Fig. 7E’, F’). This indicates that in spite of the disorganized trunk structures, such as lack of clear brain organization, the localized inducing signals required for palp initiation are still present (Wada et al., 1999; Wagner et al., 2014). However, the palps are complex structures involving epidermal and neural cells (Torrence and Cloney, 1983; Zeng et al., in press), so additional work would be required to determine whether complete palp structures develop.

Overall, basic cell type specification is accomplished in all types of TD embryos. This finding indicates that spatial relationships between cells are intact to the extent that the necessary segregation of cytoplasmic determinants and cell-cell interactions required for cell type specification can occur. It is also possible that regulative processes can compensate for some aspects of TD. As a general pattern, it appears that, given the conditions of our experiments, early stages through the gastrula are less disrupted than later stages (Figs. 5 & 6). Since basic cell types are established prior to or around gastrulation (Lemaire, 2009), this may explain why late appearing characteristics are present even in very abnormal TD embryos.

### Cleavage and gastrulation patterns are affected in TD embryos

At the temperatures we tested, gross morphology of cleavage and gastrulation stages is similar to normal embryos (Figs. 2 & 5), but closer examination reveals anomalous asymmetries in the embryos. At cleavage and gastrula stages, numbers and overall positions of cells are similar to controls, but the size and shape of cells varies, apparently randomly (e.g. Fig. 6B, Fig. 6D-E). These abnormalities will undoubtedly affect the details of cell-cell interactions. However, as described above, cell type specification events requiring specific cell relationships do occur, suggesting that cells are not grossly rearranged within the embryo. This robustness to cleavage anomalies may be due to regulatory processes compensating for some abnormalities in shape and position. A possible molecular mechanism for this regulatory compensation is that alternative inducers exist for some cell specification events. For example, as mentioned above, different blastomeres are capable of inducing prospective notochord cells to adopt the notochord cell type (Nakatani and Nishida, 1994).

### Post-gastrulation morphogenesis is disrupted at high temperature

While cell type specification works more or less normally at the threshold temperature, the architectural arrangement of those cells is disrupted. This is most apparent for the notochord. In severely affected TD embryos, the notochord cell type - large, vacuolated cells - is still apparent, but the cells fail to develop into the gently curved normal structure.

Notochord morphogenesis is generally broken down into phases (Denker and Jiang, 2012; Dong et al., 2009; Jiang and Smith, 2007; Munro et al., 2006). Initially 4 prospective notochord blastomeres divide to form a monolayer plate of 40 cells. These cells then undergo convergent extension by intercalation, resulting in a single file of cells resembling a “stack of coins”. In TD embryos the intercalation movements generally occur, although not in all cases (eg. arrow in Fig. 7D’). In normal morphogenesis the next phase is lengthening of the notochord by constriction of notochord cell diameter, driven by an actomyosin contractile ring, in combination with increase in cell volume by vacuolation. In the TD embryos vacuolation occurs, and to some extent cell lengthening, but the cells fail to maintain the gently curved configuration, and become kinked or completely disorganized. Thus, in TD embryos, notochord cells are specified and undergo the basic steps in notochord morphogenesis, but cell shapes are abnormal resulting in a kinked or twisted final configuration. One explanation for these results would be that notochord formation is largely autonomous, in combination with neural and muscle cells (Keys et al., 2002; Munro and Odell, 2002), but abnormalities in the cytoskeleton and/or the extracellular sheath that normally surrounds the notochord cause the shape defects (Jiang and Smith, 2007; Wei et al., 2017).

Along with the defects in notochord morphogenesis the other tissues of the tail, muscle, nerve cord, endodermal strand, and epidermis are also abnormal. These tissues take on the shape of the malformed notochord, reflecting their dependence on the notochord for structural support (Di Gregorio et al., 2002). Subsequent work examining notochord development in normal and TD conditions in real time might unravel the roles of these interacting tissues and morphogenetic players, and what the limiting factors are.

In terms of trunk morphogenesis, the observations show progressively more disruption of the anatomical organization of the nervous system, pharynx, and endoderm. In Type II embryos, the stomodeum and sensory vesicle are similar to controls. Type III and IV embryos do not have clearly discernable stomodaea or sensory vesicles. Obvious consistent patterns to the TD trunk phenotype were not observed, suggesting stochastic misalignments by the gastrula stage become amplified as morphogenesis proceeds. Fluorescently labeled markers followed in live embryos would help to characterize how these trunk abnormalities come about.

### Developmental mechanisms affected by high temperature and canalization

There are a number of candidate mechanisms whose breakdown could account for the disruption of morphogenesis. Many fall under the general heading of cell stress. Protein folding anomalies, redox stress (e.g. production of reactive oxygen species), and lipid damage, could all affect mechanisms of morphogenesis. For example, cytoskeletal proteins are regulated by redox signaling and susceptible to redox imbalances (Wilson et al., 2016; Wilson and González-Billault, 2015). Damage to the actin/tubulin cytoskeleton might affect the polarization and shape of cells in the nascent notochord, in turn causing defects such as those seen in TD embryos. Consistent with this notion is the finding that changes to cytoskeletal protein expression are among the most common in a proteomic survey of *C. robusta* subjected to heat shock conditions (Serafini et al., 2011; Tomanek, 2015). Definitive explanations of the mechanisms of TD effects will depend on more detailed examination of TD defects at a cellular level.

Other direct or indirect effects of temperature might be revealed by high-throughput RNA-seq at the tissue or single cell level. This kind of assay would show the extent to which transcriptional regulation is disrupted by temperature, either through misallocation of cytoplasmic factors during cell division, or disruption of normal cell-cell signaling. It is likely that full characterization of TD will depend on a combination of transcriptomic, proteomic, and cytoanatomical approaches.

A larger implication of the characteristics of embryogenesis in *C. intestinalis*, and the limits to its canalization, is that of the role of basic differences in developmental mechanisms between animal groups and their tolerance of environmental variation. For example, do highly regulative embryos, such as sea urchins, have different mechanisms limiting their canalized developmental range than more “mosaic” embryos, such as *C. intestinalis*? Detailed examination of embryogenesis within and outside the range for normal development in other species may help to answer this question. These approaches may also help to illuminate a related evolutionary issue - which developmental mechanisms are the targets of natural selection by environmental factors.

## Acknowledgements

The authors thank the Point Judith Marina for allowing collection of *C. intestinalis* from their docks. We also thank Lionel Christiaen and Thomas Meedel for perceptive comments on the manuscript, and Amaia Aldazabel for assisting in preliminary experiments.

## Funding

This work was supported by a Project Development grant from Rhode Island Sea Grant, and a Project Completion Grant from the University of Rhode Island. This material is also based upon work conducted at Rhode Island NSF EPSCoR research facilities: the Genomics and Sequencing Center and Marine Life Sciences Center, supported in part by the National Science Foundation EPSCoR Cooperative Agreement #OIA-1655221.

